# Is removal of weak connections necessary for graph-theoretical analysis of dense weighted structural connectomes?

**DOI:** 10.1101/531350

**Authors:** Oren Civier, Robert Elton Smith, Chun-Hung Yehb, Alan Connelly, Fernando Calamante

**Author notes:** Corresponding author: Oren Civier.

## Abstract

Recent advances in diffusion MRI tractography permit the generation of dense weighted structural connectomes that offer greater insight into brain organization. However, these efforts are hampered by the lack of consensus on how to extract topological measures from the resulting graphs. Here we evaluate the common practice of removing the graphs’ weak connections, which is primarily intended to eliminate spurious connections and emphasize strong connections. Because this processing step requires arbitrary or heuristic-based choices (e.g., setting a threshold level below which connections are removed), and such choices might complicate statistical analysis and inter-study comparisons, in this work we test whether removing weak connections is indeed necessary. To this end, we systematically evaluated the effect of removing weak connections on a range of popular graph-theoretical metrics. Specifically, we investigated if (and at what extent) removal of weak connections introduces a statistically significant difference between two otherwise equal groups of healthy subjects when only applied to one of the groups. Using data from the Human Connectome Project, we found that removal of weak connections had no statistical effect even when removing the weakest

~70-90% connections. Removing yet a larger extent of weak connections, thus reducing connectivity density even further, did produce a predictably significant effect. However, metric values became sensitive to the exact connectivity density, which has ramifications regarding the stability of the statistical analysis. This pattern persisted whether connections were removed by connection strength threshold or connectivity density, and for connectomes generated using parcellations at different resolutions. Finally, we showed that the same pattern also applies for data from a clinical-grade MRI scanner. In conclusion, our analysis revealed that removing weak connections is not necessary for graph-theoretical analysis of dense weighted connectomes. Because removal of weak connections provides no practical utility to offset the undesirable requirement for arbitrary or heuristic-based choices, we recommend that this step is avoided in future studies.

**Declarations of interest:** none.

## 1. Introduction

Brain connectomics focuses on obtaining a comprehensive graph of the connections in the brain (the so-called ‘connectome’, Sporns et al., 2005). One method to infer such a graph is to use diffusion MRI fibre-tracking to estimate the macroscopic axon fibre bundles that interconnect brain regions-of-interest defined typically by brain parcellations. Each reconstructed fibre track is referred to as a streamline, and from a set of streamlines (tractogram), it is possible to compute a structural connectome (e.g., Hagmann et al., 2008). The exact methods used to compute the graph from diffusion MRI streamlines tractograms can have a large influence on connectome characteristics, and the effects of a number of methodological aspects have been examined previously (de Reus and van den Heuvel, 2013; Smith et al., 2015a; van Wijk et al., 2010; Yeh et al., 2016). This article focuses on an important step in the analysis process, where one needs to choose which of the connectome’s connections to retain before performing graph-theoretical analysis.

Because traditional fibre-tracking methods cannot reliably quantify the strengths of connections in the brain (Jones et al., 2013), many studies have opted to discard connection strengths altogether and run graph-theoretical analysis on ***binary*** connectomes. The simplest method to compute a binary connectome is to generate a weighted connectome (where each edge is often coded by the number of streamlines connecting the relevant pair of regions-of-interest) and then binarize it, i.e. setting edges that have at least one streamline to “1”, and edges with no streamlines to “0”. However, if binary connectomes are computed in this simple way, that would cause weak connections to be indistinguishable from strong connections often backed up by several orders of magnitude more streamlines, thereby completely losing the putative biological heterogeneity of fibre connection strengths (Markov et al., 2011; Ypma and Bullmore, 2016). The weak connections (which are often assumed to be spurious, i.e. false positives) would then influence the subsequent graph analysis to an extent that may be undesirable (Zalesky et al., 2016). To circumnavigate this problem, the binarization of the connectome is usually preceded by the removal of weak connections^1^ (see Fornito et al., 2013; Hagmann et al.; Sotiropoulos and Zalesky, 2017; Vasa et al., 2018). These two steps combined are referred to in the literature as “thresholding”. While effective in solving the problem of weak (possibly spurious) connections having too much influence, removal of weak connections introduces its own confounds; in particular, it requires one or more arbitrary or heuristic-based choices (e.g., connection strength threshold).

Advanced probabilistic tractography methods that take advantage of voxel-level modelling that resolves crossing fibres (Tournier et al., 2011) can be combined with quantitative tractography methods (Christiaens et al., 2015; Daducci et al., 2014; Girard et al., 2017; Lemkaddem et al., 2014; Pestilli et al., 2014; Reisert et al., 2011; Sherbondy et al., 2010; Smith et al., 2013) to enable the computation of dense weighted connectomes. Such connectomes can quantify connection strength more reliably (Smith et al., 2015a), and thus, this information can be preserved and directly used for graph-theoretical analysis (binarization is not necessary). In principle, this approach should provide a deeper understanding of the brain as a network by reflecting the large range of connection strengths (Bassett and Bullmore, 2016) rather than simply discarding this information (Rubinov and Sporns, 2011); not surprisingly, dense weighted connectomes are increasingly being adopted by the research community (e.g., Conti et al., 2017; Oxtoby et al., 2017). As their name implies, dense weighted connectomes also include many more potential connections in the brain (i.e. high connectivity density). This is an advantage compared with binary connectomes which are sparse in nature; unlike weighted graphs, a fully or almost-fully connected binary graph simply does not encode any meaningful information on topology.

It might be expected that a shift from binary connectomes to dense weighted connectomes would render obsolete the problem of the over-influential weak connections, specifically so because the binarization step is not utilized anymore. Perhaps surprisingly, however, the same strategy of removing weak connections is still frequently applied also to dense weighted connectomes; a practice which we will refer to as *pruning* (for a recent example, refer to Kamagata et al., 2018; see Rubinov and Sporns, 2010). Because pruning may have substantial effects on subsequent graph analysis (Drakesmith et al., 2015; Fornito et al., 2013), we seek here to thoroughly investigate its effects on dense weighted connectomes, thus providing much-required guidelines for optimal processing of this class of connectomes. Without a standard approach to pruning in such connectomes, comparable studies might arrive at different conclusions due specifically to differences in methodology. Worse yet, it is possible that in dense weighted connectomes, pruning might actually be *disadvantageous*; if so, then studies that utilize this strategy may be unnecessarily compromised. For this reason, the current study not only evaluates the benefits of pruning, but also its potential drawbacks.

In general, pruning has some fundamental disadvantages, and thus should be carefully considered before use. From a methodological perspective, the method with fewest assumptions should be preferable; this may therefore advocate *omission* of the pruning step entirely, if it requires arbitrary or heuristic-based design choices and/or parameters yet is not beneficial to the analysis (Fornito et al., 2013; van Wijk et al., 2010). In order to perform pruning, one must indeed choose the pruning *method* – for instance, (a) subject-wise by connection strength threshold, by removing all connections up to a given connection strength; (b) subject-wise by connectivity density, by removing enough of the graph’s weakest connections such that it is reduced to a given connectivity density; or

(c) group-wise (as explored in Perry et al., 2015) -- as well as any *parameters* requisite for that method (see Rubinov and Sporns, 2010). It is of note that some studies perform their statistical analysis using several pruning parameter values in an attempt to alleviate this issue, but this approach has its own limitations (see Drakesmith et al., 2015; Fornito et al., 2013; Rubinov and Sporns, 2011).

The present work evaluates the effects of pruning on graph-theoretical metrics of dense weighted connectomes. Our goal is to critically assess whether this processing step entails enough advantages to graph-theoretical analysis of this class of connectomes to outweigh any possible drawbacks, the aforementioned and others. Because the analysis of weighted connectomes gives more weight to strong connections than to weak ones (importantly, there is no binarization step to equate them), the influence of weak connections might be reduced *intrinsically*. Thus, in practice, pruning might be redundant (see Bassett and Bullmore, 2016). To the best of our knowledge, however, this theoretical consideration has not yet been explicitly tested in dense weighted connectomes. Here we explore the effects of several pruning methods (using a range of pruning extents for each method) on selected graph-theoretical metrics derived from empirical dense weighted connectome data. Based on our observations, we then make recommendations for best practise.^2^

## 2. Methods

We analysed 100 pre-processed adult datasets from the Human Connectome Project (HCP) (Van Essen et al., 2013), which were acquired on a customized Siemens Magnetom Skyra 3T MRI system using a multi-band pulse sequence (Feinberg et al., 2010; Moeller et al., 2010; Setsompop et al., 2012; Xu et al., 2012). The diffusion imaging protocol consisted of 3 diffusion-weighted shells (b = 1000, 2000, and 3000 s/mm^2^), 90 diffusion-weighted volumes each, and 18 reference volumes (b = 0 s/mm^2^). For distortion correction, all images were additionally acquired with reversed phase encoding (Andersson et al., 2003). Other diffusion-weighted imaging parameters were: 145 x 145 matrix, 174 slices, 1.25 x 1.25 x 1.25 mm^3^ voxel size, TR/TE = 5520/89.5 ms. The HCP pipeline incorporates image reconstruction with SENSE1 multi-channel (Sotiropoulos et al., 2013), and diffusion imaging distortion correction (Andersson and Sotiropoulos, 2015, 2016). The HCP data also include high resolution T1 anatomical images, acquired using the 3D magnetization-prepared rapid gradient echo sequence (MPRAGE) (Mugler III and Brookeman, 1990) with 0.7 x 0.7 x 0.7 mm^3^ voxel size, TR/TE = 2400/2.14 ms, and flip angle = 8°.

We also analysed 10 healthy adult datasets that we acquired on an in-house, clinical-grade scanner (Siemens Skyra 3T MRI system). Informed consent was obtained from each participant prior to the study. The diffusion imaging protocol consisted of 64 diffusion-weighted volumes (b = 3000 s/mm^2^) and one b = 0 s/mm2 volume acquired using a twice-refocused spin-echo echo-planar imaging sequence (Reese et al., 2003). For distortion correction, two b=0 volumes with opposite phase encoding were also acquired (Andersson et al., 2003). Other diffusion-weighted imaging parameters were: 96 x 96 matrix, 60 slices, 2.5 x 2.5 x 2.5 mm^3^ voxel size, 8200 or 8400 ms TR, and 110 ms TE. High resolution T1 anatomical images were acquired as well, using a 3D MPRAGE sequence with 0.9 x 0.9 x 0.9 mm^3^ voxel size (256 x 256 matrix, 192 slices, TR/TE = 1900/2.64 ms).

Separate processing was performed on the HCP and clinical-grade diffusion MRI data. Where not stated explicitly, processing was performed using the *MRtrix3* software package (www.mrtrix.org). To further process the “minimally pre-processed” HCP data, we first applied bias-field correction (Tustison et al., 2010). This was followed by multi-shell multi-tissue constrained spherical deconvolution (CSD) (Tournier et al., 2007; Tournier et al., 2004) to model white matter, grey matter and cerebrospinal fluid (Jeurissen et al., 2014), with a maximum spherical harmonic degree L_max_ = 8. The diffusion MRI data from the clinical-grade scanner went through our entire pre-processing pipeline that includes denoising (Veraart et al., 2016a; Veraart et al., 2016b), Gibbs ringing removal (Kellner et al., 2016), geometric distortion correction (Andersson and Sotiropoulos, 2016; Smith et al., 2004), as well as the bias-field correction mentioned above. We then applied the multi-shell multi-tissue CSD algorithm to model white matter and cerebrospinal fluid with the same setting used for the HCP data (L_max_ = 8).

Following the initial processing, connectomes were generated in an identical fashion for both datasets using *MRtrix3*. For each subject, tractogram construction included several steps: generation of 10 million probabilistic streamlines using the 2^nd^-order Integration over Fibre Orientation Distributions algorithm (iFOD2, Tournier et al., 2010) and anatomically-constrained tractography (ACT) (Smith et al., 2012), with dynamic seeding (Smith et al., 2015b), FOD amplitude threshold 0.06, step size which is half of voxel size (i.e. step size of 0.625 mm for HCP data, or 1.25 mm for clinical-grade data), length of 5-300 mm, and backtracking, i.e. possibility for tracks to be truncated and re-tracked if a poor structural termination is encountered (Smith et al., 2012). To make quantification of connectivity biologically meaningful (Smith et al., 2015a), each streamline was assigned a weight, computed using SIFT2 (spherical-deconvolution informed filtering of tractograms, see Smith et al., 2015b). Based on each subject’s tractogram, an individual connectome was computed using 84 regions-of-interest (‘nodes’) parcellated in native space (cortex and cerebellum using freesurfer, Desikan et al., 2006; subcortical regions using FIRST, Patenaude et al., 2011, see Smith, 2015a), with connection strengths calculated by summing the weights of the relevant streamlines. Intra-node connections, which accounted for up to half of the total connection strength in the graph, were excluded before further analysis (Rubinov and Sporns, 2010).

The graph-theoretical metrics (weighted versions) interrogated in this study included: global efficiency, betweenness centrality, characteristic path length and modularity (Q-index) (Rubinov and Sporns, 2010), calculated using the Brain Connectivity Toolbox (Rubinov and Sporns, 2010); vulnerability (Iturria-Medina et al., 2008) and clustering coefficient (Zhang and Horvath, 2005). Vulnerability and clustering coefficient metrics, as well as subsequent statistical analyses, were calculated using in-house Matlab routines (The Mathworks, Inc., Natick, MA., USA). The clustering coefficient variant proposed by Zhang and Horvath was used, as a previous study found it to be more appropriate for dense weighted connectomes (Yeh et al., 2016) compared to the commonly-used version (Onnela et al., 2005).

To examine the contribution of weak connections to these metrics, we analysed each individual’s connectome separately. At each instance, we removed from the given connectome all weak connections up to a certain extent, keeping only the *X*% strongest connections of the graph (from all possible graph connections). *X%*, which is the connectivity density of the resulting pruned connectome (will be referred to as *target density*), ranged from the density of the original unmodified connectome (no pruning), and down to 0% density (all connections pruned). At each target density *X%*, we then quantified a range of key weighted graph-theoretical metrics. Moreover, we converted each metric from its absolute value to a relative change from the *baseline value* (i.e. the value for that metric as calculated on the original unmodified connectome).

Our initial *qualitative* evaluation simply involved plotting, for each metric and for each subject, the relative change from baseline (as defined above) when the subject’s connectome is pruned to different target densities. We also examined relative changes in the group mean metric values (separately for each dataset). To calculate the group mean value for a specific target density, we pruned all individual connectomes to that density, calculated the graph-theoretical metric for each individual, and averaged these values across subjects. As the original graph density varies across individuals, if not all individual connectomes of a dataset could be pruned to a particular target density, no group mean was calculated for that density. For example, because the HCP cohort includes a subject whose connectome had a baseline density of 78%, we calculated the mean values of the HCP dataset for all densities ≤78%.

To investigate when pruning is consequential in the *statistical* sense, we tested which extents of pruning can significantly differentiate between two groups of healthy adult connectomes, i.e. generate a statistical difference between the groups when applied to only one of them. For this experiment, we drew two groups of subjects from our 100-subject HCP cohort by randomly splitting it in half. We repeated the experiment 1000 times to ensure the results are not dependent on a specific split. The complete analysis was performed as follows, separately for each metric:

1. For each of 1,000 repetitions:
  a. Randomly split the HCP cohort into two equal groups (50 subjects in each): “pruned” and “unpruned”.
  b. For every *feasible* target density *X*% (i.e. a density where a mean can be calculated, see previous paragraph):
    1. Prune the connectomes of those subjects in the “pruned” group to the target density *X*%, but leave the connectomes of the subjects in the “unpruned” group unmodified.
    2. Calculate graph-theoretical metric value for all subjects.
    3. Generate *t-statistic* by performing two-sided t-test between metric values of the “pruned” and “unpruned” groups.
    4. Generate *null distribution* by doing the following for each of 10,000 repetitions:
      a. Randomly permute the group labels (“pruned” and “unpruned”)
      b. Perform two-sided t-test between metric values of the permuted “pruned” and “unpruned” groups.
    5. From the t-statistic calculated in 1.ii.(3), and the null distribution generated in 1.ii.(4), produce a nonparametric *p-value*.
2. For every feasible target density *X%*:
  1. Report the number of repetitions (out of the total of 1,000) for which the *p*-value calculated in 1.ii.(5) was less than 0.05.

It should be noted that due to the nature of the analysis, some splits will result in groups that are significantly different even without any pruning taking place: this is an inevitable consequence of statistical testing over large samples where small sample-specific differences might erroneously lead to the rejection of the null hypothesis (in about 5% of cases). That said, because our analysis includes many splits, the existence of these few outliers is expected to be evident in the result plots (these few splits will have significant *p*-values all along the density range), which should permit us to effectively disregard them during evaluation.

We performed two additional analyses to a subset of the HCP data (10 subjects chosen arbitrarily) to investigate the generality of the overall observations to variants of our analysis approach. The first analysis employed a different pruning method: instead of pruning individual connectomes to a certain density, we removed from individual connectomes all connections whose strengths are below a certain threshold. The second analysis employed a different parcellation scheme for connectome computation: a higher-resolution ‘Laussanne2008’ parcellation. The Laussanne2008 parcellation is based on a FreeSurfer parcellation (Desikan et al., 2006) whose regions are subsequently subdivided, such that all the resulting smaller regions are approximately equal in size (Hagmann et al., 2008). We utilized the Laussanne2008 parcellation image (234-regions version), and nonlinearly transformed it onto each of our 10 HCP subjects. This was performed using the Nipype interface (Gorgolewski et al., 2011) of the connectome mapper toolkit (cmtk; http://www.connectomics.org) (Daducci et al., 2012). As a larger number of regions in these connectomes leads to a larger number of potential inter-areal connections, for this analysis we constructed tractograms with 70 million rather than 10 million streamlines.

## 3. Results

### 3.1. Qualitative analysis

The 100 dense weighted connectomes computed for our HCP cohort had an average density of 89.9%. Fig. 1 shows the effect of pruning on the graph-theoretical metrics (weighted versions) of 10 representative individual connectomes, as well as on the group mean graph-theoretical metrics, calculated on the whole cohort of 100 connectomes. Examining the panel for the global efficiency metric from left to right (high to low target densities for pruning), it is evident that there is not much change in this metric value, even when pruning down to very low densities. If we take for example a ±5% change from the baseline value as a criterion (i.e. ±5% from the metric calculated on the original unmodified connectome), the decrease in global efficiency did not exceed −5% all the way down to about 6.5% target density. Moreover, this behaviour characterized all subjects, as evident by the strong similarity between the trajectories of the 10 representative individuals (circle in Fig. 1 shows the point of exactly −5% decrease in the group mean). Similar trends were observed in betweenness centrality and characteristic path length, albeit with somewhat greater inter-subject variability. The trend of limited change from baseline down to a low target density applied also to vulnerability, with the difference that the metric increases rather than decreases with lower densities. Another observation from Fig. 1 is that for the first four metrics in particular, once they begin to deviate from the baseline value with lower target densities, a high rate of change is reached very quickly.

**Fig. 1.**
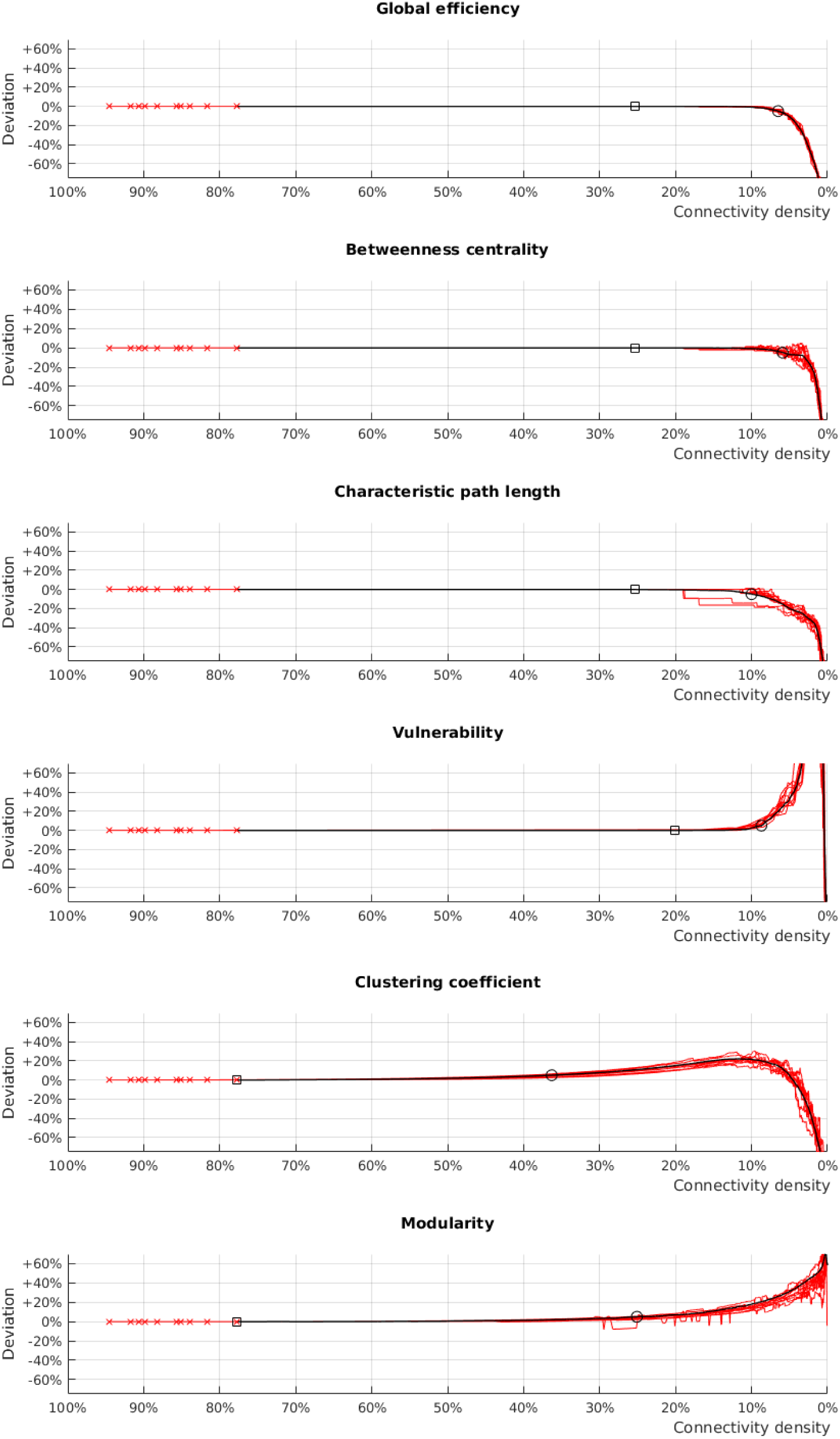
The effect of pruning on graph-theoretical metrics (weighted versions) of dense weighted structural connectomes. For each graph-theoretical metric, the relative change in the metric (compared to the original unmodified connectome) for different target densities is shown both for 10 representative subjects (red lines) and the mean of all 100 subjects (black line; see Section 2 for the calculation of the group mean). Red crosses: the connectivity densities of the original unmodified connectomes of the 10 representative individuals (i.e. no pruning applied). Squares: examining the panels from left to right, the first target density where at least one of the 100 individuals has a detectable change in metric value relative to the original unmodified connectome. Circles: examining the panels from left to right, the first target density where the change in group mean relative to the baseline group mean reaches +5% or −5%.

In contrast with the first four metrics in Fig. 1, in the lower two metrics (clustering coefficient and modularity), changes in the metrics due to pruning commenced at higher densities, and were also more gradual. Using the same criterion as above, the point of exactly ±5% change in the group mean modularity value is at 25.1% target density. Down from this target density, the modularity metric initially changes gradually, but with the change rate accelerating as density approaches 0%. The case of clustering coefficient is unusual: exactly +5% change in group mean occurs at 36.3% target density (the highest figure across all metrics studied), but at about 10% density, the metric turns from gradually increasing with lower density, to rapidly decreasing with it (the vulnerability and modularity metrics exhibited a comparable effect, but the switch in the direction of change occurred there at extremely low densities that are not meaningful to graph-theoretical analysis of connectomes).

The analysis of global efficiency also showed that pruning did not introduce any detectable change, not even in a single subject, down to a target density of 25% (Fig. 1, top, square; changes are detected at the default precision provided by Matlab, double-precision floating-point). Moreover, for most of the individuals, global efficiency was resistant to change down to even lower target densities (range of densities for the group 11%-25%, median 14%; the small changes, where they occurred, are too small to be observed in the figure). This trend of complete insensitivity to pruning across most of the target density range was consistent across the first four metrics. Fig. 2 demonstrates these findings qualitatively: pruning one of the connectomes from a baseline connectivity density of 93% to a density of 31% (i.e. removing two thirds of the connections) revealed no detectable change from baseline value in any of the first four metrics examined.

**Fig. 2.**
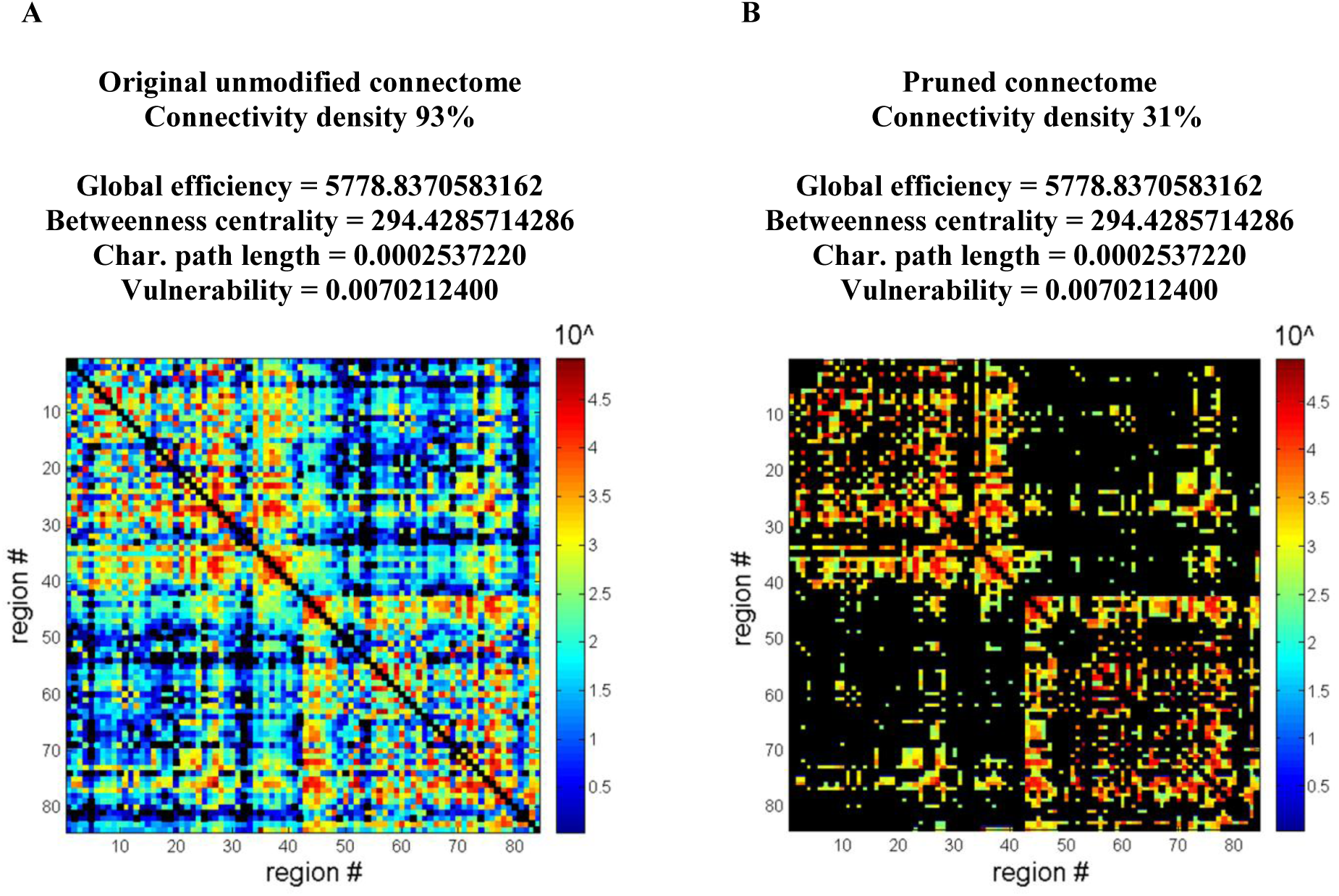
Pruning produces no detectable change from baseline in four key graph-theoretical metrics (weighted versions) down to relatively low densities. In this example, (A) shows the original unmodified connectome of one of the subjects from Fig. 1, and (B) shows the same connectome pruned down to 31% density. It is of note that despite removing two thirds of existing connections, there was only a minor effect on the sum of connections strengths in the graph, which was reduced from 5,122,728 at baseline to 5,005,006 after pruning. The change is minor because the connections removed were relatively weak (for this connectome, a 31% target density corresponded to a connection strength threshold of 273). Colorbar shows connection strength. Missing/removed connections are in black.

### 3.2. Quantitative statistical analysis

To quantitatively investigate when pruning is consequential, we tested which pruning extents can generate a statistical difference between two groups of otherwise similar connectomes when applied to only one of the groups. Fig. 3A demonstrates this analysis for a representative group split of our HCP cohort; shown are the graph-theoretical metric means and standard deviations (SDs) (for the “pruned” and “unpruned” groups) that were used to calculate a *p*-value for each target density analysed. We repeated the analysis on 1000 random splits of the original cohort, and Fig. 3B shows the *p*-values as a function of target density for 20 of these splits (each line represents an individual split).^3^ Fig. 3C summarizes the results of these statistical tests by showing the number of splits (out of 1000) that resulted in a significant group effect (*p* < 0.05) for each target density analysed.

**Fig. 3.**
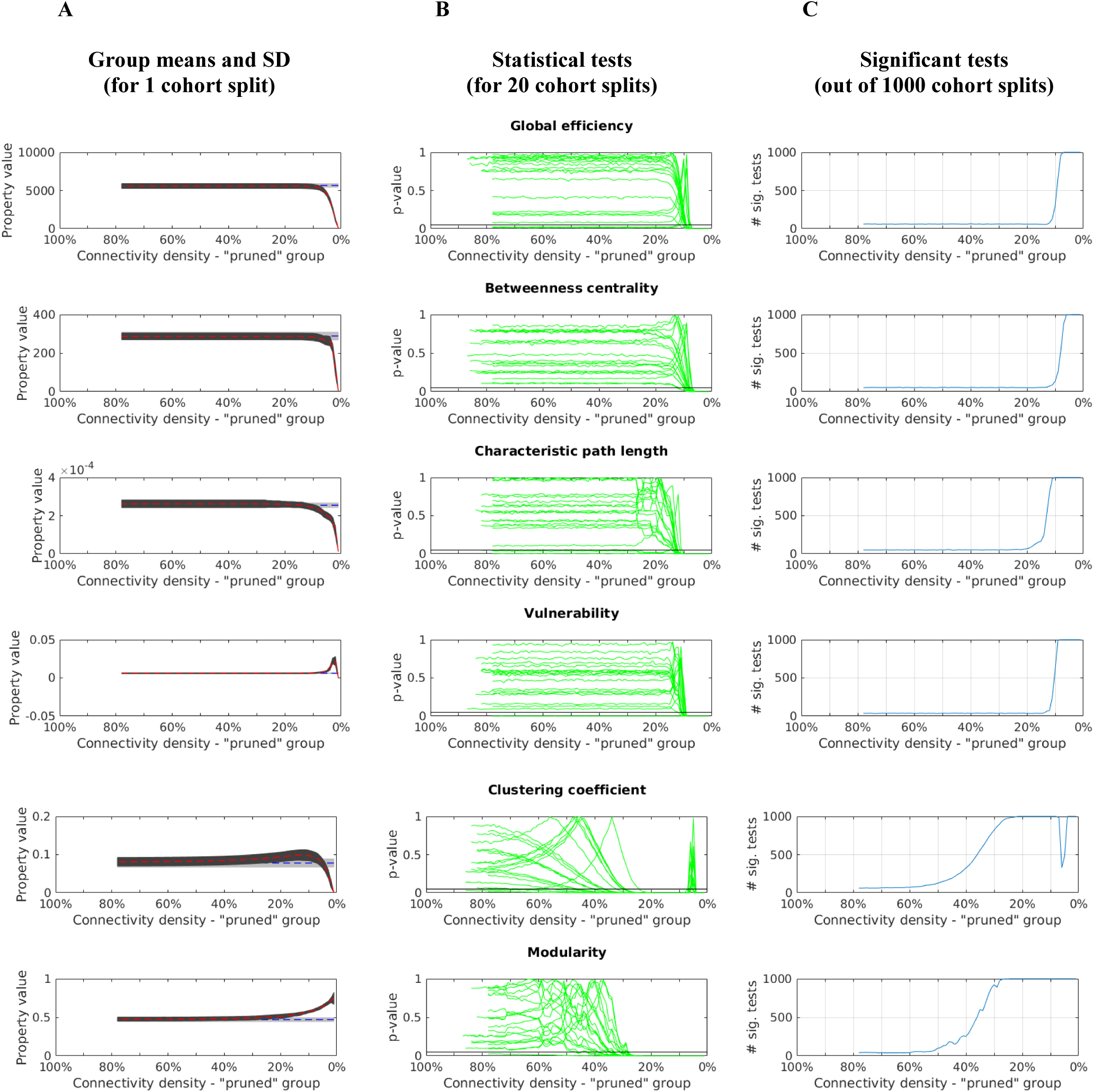
Statistical analysis of the effect of pruning on graph-theoretical metrics (weighted versions) calculated in dense weighted connectomes. (A) Means (dashed lines) and standard deviations (SD, dark and light grey bands) for graph-theoretical metrics of “pruned” (red) and “unpruned” (blue) groups of connectomes for a representative split of the original cohort. Data are shown for the range of feasible target densities (see Section 2). The means and SDs of the “unpruned” group are constant, thus shown as flat lines and bands. (B) p-values for the comparison between the graph-theoretical metrics of “pruned” and “unpruned” groups; for visual clarity, results are shown for only 20 out of the 1000 random splits. The horizontal black line at the bottom of each panel marks p = 0.05. (C) For each target density, the number of splits out of 1000 where the “pruned” and “unpruned” groups had significantly different (p < 0.05) graph-theoretical metrics.

The results shown in Fig. 3C address the fundamental question of “Does pruning have an effect?”, by demonstrating the likelihood of detecting a significant change from baseline at *p* < 0.05 due to the application of a pruning step. When applying very little pruning or a lot of pruning, the answer is the same for all metrics. At very little pruning (target density > 50%), a significant change from baseline is detected in only around 50 of the 1000 splits. This does not support an underlying effect, as at least ~5% rejections of the null hypothesis are expected to always occur when performing many statistical tests, whether an effect exists or not (see Section 2). Conversely, applying a lot of pruning (target density < 10%) does have an effect as it introduces a statistically significant group difference across all metrics investigated, regardless of how the subject data are split randomly into “pruned” and “unpruned” groups. This is evident by a significant group difference being detected in all 1000 splits. The transition between these two behaviours however does differ slightly between the metrics. For the first four metrics in Fig. 3 (global efficiency, betweenness centrality, characteristic path length, and vulnerability), this transition in the effect of pruning occurs relatively sharply at a target density of 10-15%. For the last two metrics (clustering coefficient and modularity), on the other hand, it is not predictable whether or not pruning will have an effect within the wide target density range of ~20-60%. In that range, the number of splits where a significant group difference is detected is anywhere from ~50 and above.

The results of repeating the main experiment with modifications to the experimental design are shown in Supplementary materials: Supplementary material S1, using clinical-grade data acquired in-house; Supplementary material S2, using an alternative pruning method based on connection strength threshold; Supplementary material S3, using a higher resolution parcellation scheme, with smaller, equally-sized regions. Overall, the results from these additional analyses replicate the general findings reported in the main text, showing that our observations should be generalizable to other connectomics experiments if similar tractography techniques are used for connectome generation. It should be noted however that despite the overall agreement, there are minor differences in some of the analyses included in Supplementary material S3. They will be addressed at the relevant point in the discussion (Section 4.1.4).

## 4. Discussion

The goal of this study was to critically evaluate the utility of pruning in the analysis of dense weighted structural connectomes. To this end, we examined the effects of different pruning extents applied to connectomes of healthy adult individuals. Specifically, we examined how pruning to different connectivity densities modifies key (weighted) graph-theoretical metrics, and the behaviour of the metrics as a function of the target density.

### 4.1. Empirical evaluation

As will be further discussed below, our empirical analyses revealed that, in general, there is no extent of pruning that is beneficial for dense weighted connectomes. As the effects of pruning varied with target density (including any potentially detrimental effects), we will separate the discussion into two gross pruning levels: *light-to-moderate pruning* where the pruning step is *not* consequential (the part of the density range from high target densities to densities as low as 30%, or even 10% for some metrics), and *heavy pruning* where the pruning step is consequential, but also has ramifications regarding the stability of the statistical analysis (the rest of the density range).

#### 4.1.1. Light-to-moderate pruning

In computation of binary connectomes, removal of weak connections, for example, through thresholding using a connection strength threshold greater than 0, is primarily required to solve the problem of the over-influential weak connections (van Wijk et al., 2010; Zalesky et al., 2016). In weighted connectomes this problem might be obsolete, but because many weak connections are probably still spurious, it is presumed that the removal of weak connections is required nonetheless. This apparent presumption is however contradictory to our results. We showed that unlike binary connectomes, in dense weighted connectomes removal of weak-to-moderate strength connections is actually non-consequential: pruning to densities as low as 10-30% (depending on the particular metric) did not produce any significant effect on the graph-theoretical metrics studied here. If light-to-moderate pruning is non-consequential, it obviously cannot be a solution for any underlying problem such as spurious connections.

This demonstrates that weak-to-moderate strength connections in dense weighted connectomes in practice make a negligible contribution to the calculation of the weighted graph-theoretical metrics studied here. In this class of connectomes, therefore, the influence of spurious weak connections is effectively minimized even without any pruning performed (and a by-product of this, of course, is that also real weak connections weigh less in metric calculation). This behaviour is a direct outcome of two factors combined. First, the contribution of individual connections to weighted metrics being directly dependent on the connections’ strength. And second, the long/heavy tail connection-strength distribution across many orders of magnitude observed in the connectomes analysed here and in previous studies (Smith et al., 2015a).

The per-edge connection-strength distribution that characterize our connectomes is presented in Fig. 4 and Table 1. It is striking that the weaker 80% connections of the graph are concentrated in a very small range of low connection strengths (~ 0-1,000), whereas the 20% strongest connections are spread over a large range of much stronger strengths (~ 1,000-90,000). Within this extreme distribution, most connections are very weak compared to the strongest connections in the graph, and their contribution to global metric values is accordingly very small. It is important to note that such a connection-strength distribution was demonstrated in brains across species (Ercsey-Ravasz et al., 2013; Oh et al., 2014), and thus is expected to characterize connectomes generated with other algorithms as well, as long as they adequately capture the underlying biology.

**Fig. 4.**
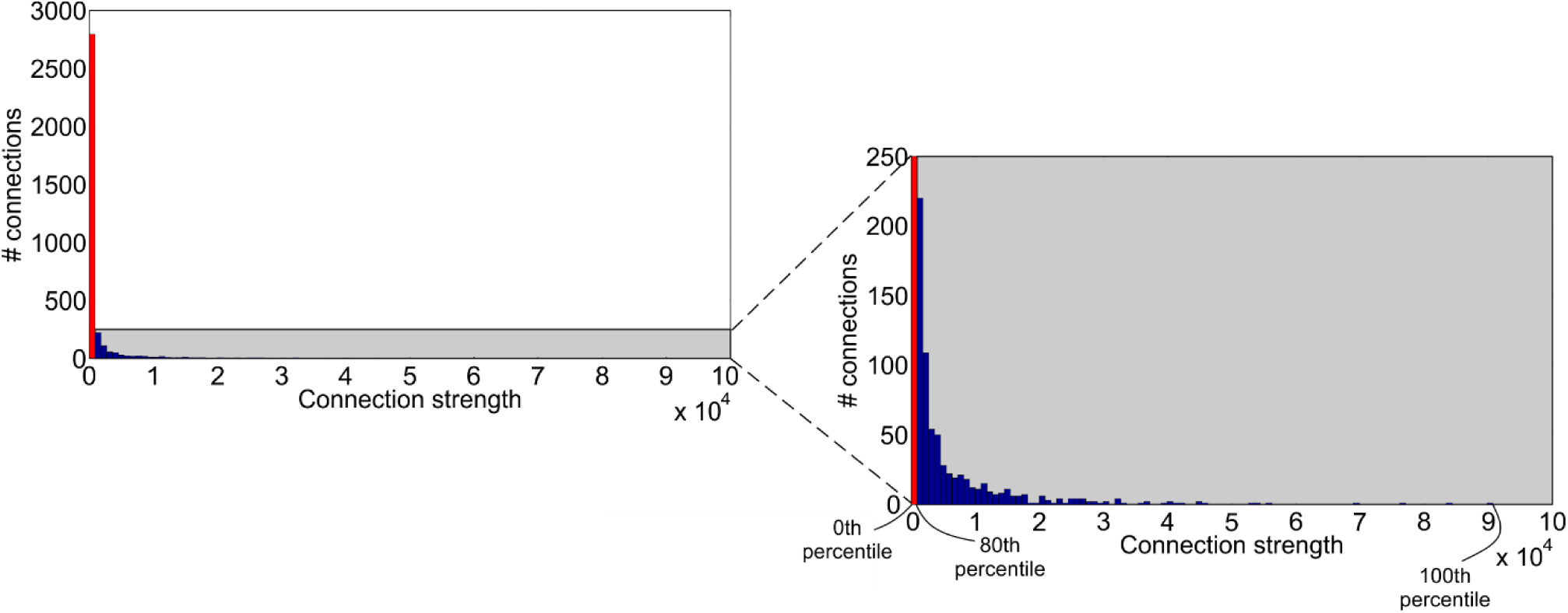
A histogram of edge-wise connection strengths in a representative HCP subject (the same subject from Fig. 2). It shows that strong connections have strengths of up to ~90,000, whereas most connections of the graph, marked in red (~80% of connections), have strengths of no more than ~1,000. The right panel is a zoomed version of the shaded region in the left panel. See also Table 1 for the exact values at different percentiles of the connection-strength distribution.

**Table 1.** Values at different percentiles of the connection-strength distribution shown in Fig. 4.

| Percentile | 0<sup>th</sup> | 10<sup>th</sup> | 20<sup>th</sup> | 30<sup>th</sup> | 40<sup>th</sup> | 50<sup>th</sup> | 60<sup>th</sup> | 70<sup>th</sup> | 80<sup>th</sup> | 90<sup>th</sup> | 100<sup>th</sup> |
| --- | --- | --- | --- | --- | --- | --- | --- | --- | --- | --- | --- |
| Connection strength | 0 | 1 | 4 | 12 | 27 | 62 | 137 | 298 | 881 | 3009 | 90710 |

In light of the above discussion, it becomes clear why the behaviour of binary connectomes is radically different from that of weighted connectomes. The binarization step applied in the computation of binary connectomes wipes out the connection-strength distribution of the original weighted graph (see Section 1), making all connections equal in strength. The connections that were originally weak have then the capacity to contribute just as much as even the strongest connections of the brain. If removal of weak connections is not applied, this can have considerable consequences (Fornito et al., 2013), and especially so if the original graph included spurious weak connections (Zalesky et al., 2016).

#### 4.1.2. Heavy pruning

The topology of the strongest connections in the connectome is considered to be more important for global brain organization than the topology of the rest of the connectome’s connections (Drakesmith et al., 2015; Rubinov and Sporns, 2010). Heavy pruning removes *all but* the strongest connections of the connectome, which is supposed to eliminate confounds that somehow masks/reduces statistical effects relating exclusively to the topology of the strongest connections (Fornito et al., 2013; van Wijk et al., 2010).

Examining our results, it is evident that when pruning to low enough target densities (<10% or even just <30%, depending on the metric), the removal of weak connections is indeed consequential. Therefore, unlike for light-to-moderate pruning, we cannot unequivocally reject the option that heavy pruning is overall beneficial, and a more detailed analysis is warranted. Below, we suggest mandatory criteria for pruning to be justifiable, and evaluate heavy pruning against them.

#### 4.1.3. Criteria to justify pruning

We suggest two mandatory criteria to evaluate whether it is justifiable to prune a connectome from its original density to a target density:

1. The pruning must lead to a predictably significant effect on the values of the graph-theoretical metric; if it does not, then the pruning step serves no purpose.
2. Following pruning, the value of the metric should not be density-sensitive, i.e. it should not be strongly dependent on the particular target density chosen. The reasoning is as follows: If the value of the metric is density-sensitive at the target density, then the particular value of that metric for an individual subject will be dependent on two additional factors other than underlying graph topology. First, the density at which the metric begins to deviate from its value at the original unmodified connectome (a density which may vary between subjects), and second, the rate of change of that metric as a function of density (which may vary between subjects as well). In such a scenario, not only does interpretation of any observed significant result becomes exceptionally difficult, but any statistical result may be unstable with respect to the precise density chosen.

Fig. 5 provides a diagrammatic representation of a hypothetical case where these criteria are satisfied across a particular range of target densities, and contrasts it against our empirical data. The cyan lines in Fig. 5 follow the same conventions as the lines in Fig. 3C; each line shows the number of random splits of a healthy cohort into “pruned” and “unpruned” groups that exhibit statistically significant group difference. In order to satisfy criterion 1 above, the line should be at or near 1000 at the target density, indicating that a significant effect is predictable. The orange lines in Fig. 5 follow the conventions of the black lines in Fig. 1; each line shows the relative change in the group mean metric value (compared to the group mean metric value of the original unmodified connectomes). In order to satisfy criterion 2, the line should be approximately flat in the vicinity of the target density, indicating negligible sensitivity to density. The black box in Fig. 5A indicates the range of target densities where these criteria are satisfied in our hypothetical case (i.e. a “sweet zone”). Pruning to any of these densities has a predictably significant effect, and at the same time, metric value does not become density-sensitive.

**Fig. 5.**
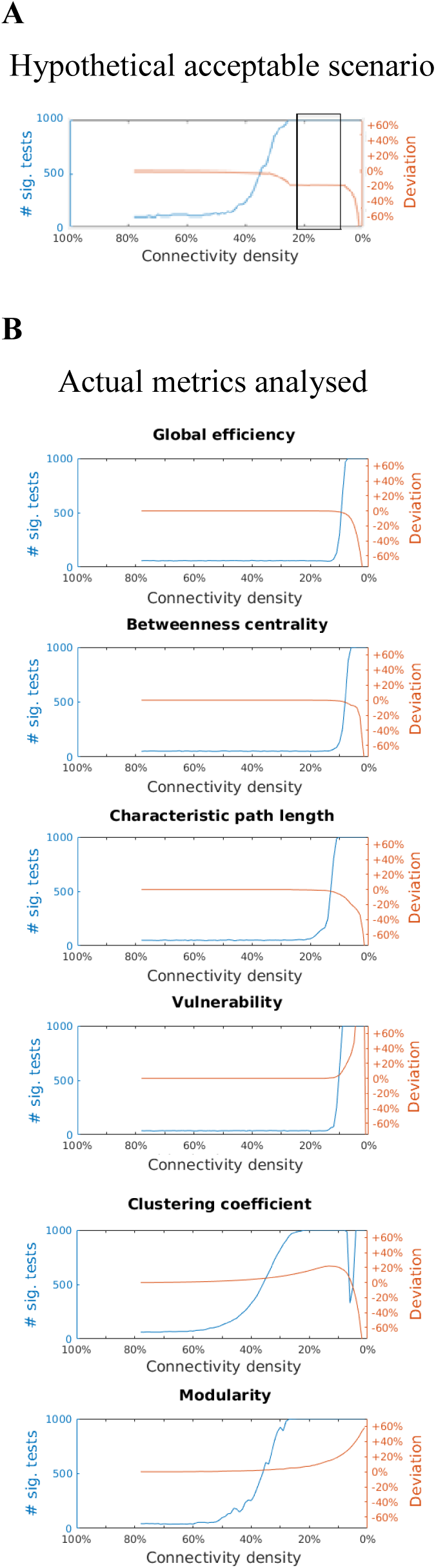
Comparison between the consequences of pruning in a hypothetical acceptable scenario and empirical data. (A) Hypothetical acceptable scenario. We have hand-drawn these trajectories to demonstrate when our two criteria for pruning would be satisfied (see Section 4.1.3). (B) Empirical behaviour of the metrics studied. Shown over all feasible target densities are: the number of cohort splits where pruning only the subjects in the “pruned” group leads to a significant group effect (cyan; see Fig. 3C for more details), and the relative change in the mean of the metric which is induced by pruning (orange; see Fig. 1 for more details). The hypothetical scenario includes a “sweet zone” (black rectangle), i.e. a range of target densities where pruning produces a predictable effect (cyan line is at the maximum of 1000), and the metric is insensible to density (orange line is flat).

Next, we evaluate what gross pruning levels satisfy both criteria. As discussed above, light-to-moderate pruning is not consequential for any metric examined and thus it does not even satisfy criterion 1. On the other hand, heavy pruning is consequential for all the metrics, hence, it does satisfy the first criterion. We shall now evaluate whether heavy pruning also satisfies criterion 2.

Examining Fig. 5B, it is evident that when heavy pruning satisfies criterion 1 (at target densities where the cyan lines are at or near 1000), criterion 2 is generally not satisfied (orange line is not approximating a flat line). This is unlike the “sweet zone” in the hypothetical case in Fig. 5A where both criteria are satisfied (cyan line at 1000 and orange line is flat). Heavy pruning thus brings the metric into a connectivity density range where it is density-sensitive, and this constitutes a potential important source of inter-subject variability as explained above. The inter-subject variability of the point where the metrics begin to deviate, and of the rate of change, can be appreciated by observing the behaviour of the metrics at that range (range of low densities) in Fig. 1.

It should be also noted that after heavy pruning of a dense weighted connectome, the resulting connectome is of course much sparser. Interestingly, studies on sparse weighted connectomes showed high sensitivity to density as well (Drakesmith et al., 2015; Yan et al., 2018). We would like emphasise, however, that our results are not replicating the previous studies but are rather new findings. The reason is that similar sparsity does not automatically imply similar sensitivity to density, and the relationship between sparsity and density-sensitivity needs to be shown for every class of connectomes separately. The previous studies investigated connectomes generated using deterministic tractography (i.e. the output of the fibre tracking algorithm is a sparse connectome), and we investigated connectomes generated using probabilistic tractography (i.e. the output of the algorithm is a dense connectome, which is then turned into a sparse connectome by means of heavy pruning).

#### 4.1.4. Evaluating special cases of pruning

Although heavy pruning does not generally satisfy both criteria 1 and 2, there is a notable exception. It occurs in the analysis of the clustering coefficient, at about 10% target density, the point along the density range where the metric gradually turns from increasing with lower density to decreasing with lower density. At that target density, the cyan line in Fig. 5B (clustering coefficient panel) is at 1000 (i.e. the effect of pruning is predictively significant), whereas the orange line is just lightly curved thus approximating a flat line (indicating low sensitivity to density). However, the orange line indicates the change in the group mean metric value and not in individual subjects. The trajectory for individual subjects actually varies (see red curves in Fig. 1, clustering coefficient panel), which in practice, makes the usefulness of this target density limited: although some subjects will have approximately flat trajectory at that point, others will not. Moreover, for some subjects the transition from increasing to decreasing with lower densities is sharper (not approximating a flat line), which would introduce even additional variability.

A behaviour similar to that observed in the clustering coefficient (shift from increasing to decreasing) is also associated with betweenness centrality and characteristic path length, calculated on the connectomes we computed using higher-resolution parcellation (supplementary material S3). However, the curvature at the point of the shift is even sharper, and thus, the reservations mentioned above apply to a greater extent.

#### 4.1.5. Wrapping it all together

We examined the effect of the whole range of pruning extents on several variants of our analysis approach (see also Supplementary Materials). The results show that only heavy pruning is consequential. However, regardless of the utility of this processing step (if any), heavy pruning also increases inter-subject variability considerably at most target densities, which may lead to unstable statistical results. This makes heavy pruning largely inadequate for dense weighted connectomes. Based on these results, we conclude that performing no pruning is the best option available.

### 4.2. Arguments for pruning that do not apply to dense weighted connectomes

Our conclusion that omitting the pruning step is the best option for dense weighted connectomes might appear contradictory to other motivations for pruning. Below we discuss why these considerations are not applicable to dense weighted connectomes.

One reason that pruning is advocated is to obtain a connectome that has small-world properties, as this is considered a fundamental characteristic of the brain across species. While binary connectomes need to be of low density to achieve small-worldness (and thus removal of all but the strongest connections is usually required), this is not the case with dense weighted connectomes, which have small-world properties despite their high density (as long as metrics appropriate for weighted graphs are used, see Bassett and Bullmore, 2016; Yeh et al., 2016). It is also argued that some graph-theoretical metrics require a minimum level of graph sparsity to be applicable (Fornito et al., 2013; Sotiropoulos and Zalesky, 2017). However, these are again criteria specific to binary connectomes, as weighted graph-theoretical metrics are designed to be applicable across the whole range of connectivity densities.

Because original individual connectomes often have different connectivity densities, the pruning step is often applied to fix connectivity density across subjects (Sotiropoulos and Zalesky, 2017). This removes possible variabilities due to the dependence of graph-theoretical metrics on connectivity density (Fornito et al., 2013; Rubinov and Sporns, 2010; van Wijk et al., 2010). However, in contrast with the sensitivity to density we indeed observed at low target densities (see Section 4.1.3), graph-theoretical metrics are completely insensitive to density at the high connectivity densities of the original connectomes (no difference between subjects at the left-hand side of Fig. 1). Moreover, even if the average density across subjects differs between two studies of the same population (e.g., if due to differences in measurement noise, more weak connections are discovered in one study than the other), that will not automatically lead to a disagreement in the results; at high densities, group-mean metrics are not sensitive to density either (Fig. 3A). We may conclude then that for dense weighted connectomes, there is no need to set a common connectivity density via pruning.

Another suggested motivation for pruning is to minimize the adverse effects on graph-theoretical metrics, both of spurious connections and missing real connections, i.e. connections that exist in the brain but are not captured by the fibre-tracking algorithm (Zalesky et al., 2016). Zalesky et al. calculated the optimal target density for such pruning to be in the range of 40-50% in binary connectomes, but no such recommendation has been made to date for weighted connectomes. It is of note, however, that if similar figures apply to weighted connectomes, extending the range of target densities up to the maximal possible density (i.e. no pruning, which is our recommended approach) would make little difference. The connections added to the connectome when increasing target density above 40% are weak (regardless of their classification into spurious or real connections), and as we showed above, their contribution to metric values is negligible.

### 4.3. Light-to-moderate pruning is generally not detrimental

It is important to note that the conclusions from our empirical analysis are two-fold – we showed that pruning as a whole is unnecessary for dense weighted connectomes, but we also showed that in the case of light-to-moderate levels of pruning, the removal of connections is not causing harm either; in fact, for some of the studies published prior to our recommendations, the level of pruning used is likely to fall into this latter category. However, the precise point where the effect of pruning shifts between not causing harm and being detrimental depends on the connection-strength distribution of the connectome studied, and this in turn relies on factors such as data acquisition and diffusion tractography techniques utilized. Therefore, we suggest that in cases where pruning is performed after all (for example, if future insights will indicate that pruning has benefits that we have not considered), the quantitative results presented here could serve as a guideline for the extent of pruning that is likely to have limited consequences in connectomes generated from HCP-quality or clinical scanner-grade MRI data, and using advanced tractography techniques.

### 4.4. Heavy pruning is detrimental

In contrast with light-to-moderate pruning, heavy pruning does have drawbacks that make it impractical in practice. Specifically, the sensitivity to the connectivity density of the resulting graph introduces inter-subject variability, and this may make any statistical result unstable and difficult to interpret. Therefore, we strongly recommend against it unless the drawbacks can be somehow mitigated.

One possible approach to overcome the drawbacks of heavy pruning is to try several pruning extents, with the goal of finding a range of extents where inter-subject variability is low enough to allow stable statistical results. This introduces however a problem of multiple-comparisons (Fornito et al., 2013; Rubinov and Sporns, 2010). The most popular solution for this newly-generated problem is to calculate the area under the curve (AUC) of the statistic across different extents of pruning. However, this solution still requires to arbitrary specify a range of pruning extents and is generally insensitive to effects limited to narrow ranges (Fornito et al., 2013). Drakesmith et al. (2015) proposed an alternative data-driven technique that identifies a range of pruning extents, within which all levels produce a statistically significant effect in the data. Using permutation correction, this technique can also decide if this detected sustained effect is larger than what would be expected by chance. This solves the multiple-comparison problem as well as the need for an arbitrary choice of parameters. While these authors showed the merit of their technique only for the case of sparse weighted connectomes, it might be also effective in the dense weighted connectomes investigated here.

Despite the advantages of the Drakesmith et al. (2015) technique, there are some potential limitations. First, Drakesmith et al. mention that interpreting significant results obtained using their technique can be challenging, for example, if the technique selects different density ranges for different metrics. This is a likely scenario in dense weighted connectomes given the range of behaviours demonstrated by weighted graph-theoretical metrics (Fig. 1). Second, it is not clear if the technique would prove sensitive enough when using pruning by connectivity density, as the paper where it is presented only explores pruning by connection strength threshold (Drakesmith et al., 2015). Pruning by connection strength threshold may raise challenges in data interpretation, and thus, is a less viable option. For example, if two groups have different distribution of connection strengths, such a pruning approach might result in different connectivity densities across the groups (Fornito et al., 2013), which leads to biases (Rubinov and Sporns, 2010; van Wijk et al., 2010). A rejection of the null hypothesis by the technique might then be a consequence of trivial differences in connectivity density, putting into question any interpretation related to topology (Fornito et al., 2013; Sotiropoulos and Zalesky, 2017). More comprehensive investigation of these points is warranted but is beyond the scope of the present study.

### 4.5. Graph-theoretical metrics that rely on shortest paths

As discussed above, light-to-moderate pruning is non-consequential for the first four metrics studied (global efficiency, betweenness centrality, characteristic path length, and vulnerability), and heavy pruning is consequential for these metrics only from target densities of about 10% and lower. Indeed, Fig. 1 shows that the values of these metrics barely change through most of the density range. Yet, a rather surprising finding from the analysis is that for part of this range (down to pruning densities of about 20%-30%) we could not detect *any* change in metric value, and that was true for all individuals (see squares in Fig. 1). This can be contrasted with the other two metrics, where changes in metric values, albeit too small to be visible in Fig. 1, were detected even for the smallest extent of pruning (indicated by the squares appearing at high target densities).

There are two factors contributing to this extreme insensitivity to pruning. The first factor is that the first four metrics (but not the last two) require the identification of the *shortest path* between each pair of regions in the graph, with metric value being calculated exclusively based on the connections in these paths (ignoring the rest of the connection in the graph). Moreover, in contrast to binary graphs, where shortest path is defined as the *smallest number of connections* that need to be visited to travel between a pair of nodes, for comparable metrics to be calculated in weighted graphs, a measure of ‘length’ must be derived for each connection, typically taken as the reciprocal of its strength (the most popular definition of connection ‘length’, which we also used here); the shortest path is then the sequence of connections with the *minimum sum of connection ‘lengths’*. Due to the use of a reciprocal, a path made of stronger connections will have a lower sum of connection ‘lengths’, and is therefore more likely to be identified as a shortest path. The biological motivation is that stronger neural connections are capable of carrying more information, and thus are more likely to provide the routes for information transfer in the brain.

The second contributing factor is the long/heavy tail connection-strength distribution reported above (see Section 4.1.1). In this distribution there are relatively few very strong connections that are several orders of magnitude greater than the majority of connections in the graph (Fig. 4 and Table 1). Yet, there are enough of these strong connections such that they can be combined to travel between each pair of regions without the need to visit any weak connections en route. Although these paths made exclusively of strong connections might include several connections each, each path will still have a total low sum of connection ‘lengths’ (as the reciprocal of the strength of each individual strong connection is extremely low).

The combination of these two factors - the conventional definition of shortest path in weighted graphs, and the characteristic connection-strength distribution of dense weighted connectomes - has a large impact: for each possible pair of brain regions, the path with the minimum sum of connection ‘lengths’, i.e. the shortest path, is one of the exclusively-strong-connections paths described above, and strikingly, this is the case even if two regions are directly connected by a weak connection (see Fig. 6). The strong connections will be then the only connections that weigh into the calculation of the four graph-theoretical metrics mentioned above (i.e. those that rely on shortest paths). This explains why these metrics are not affected by pruning of only the weaker connections of the graph: because the weak connections are effectively ‘bypassed’ during metric calculation, their removal has no effect whatsoever.

**Fig. 6.**
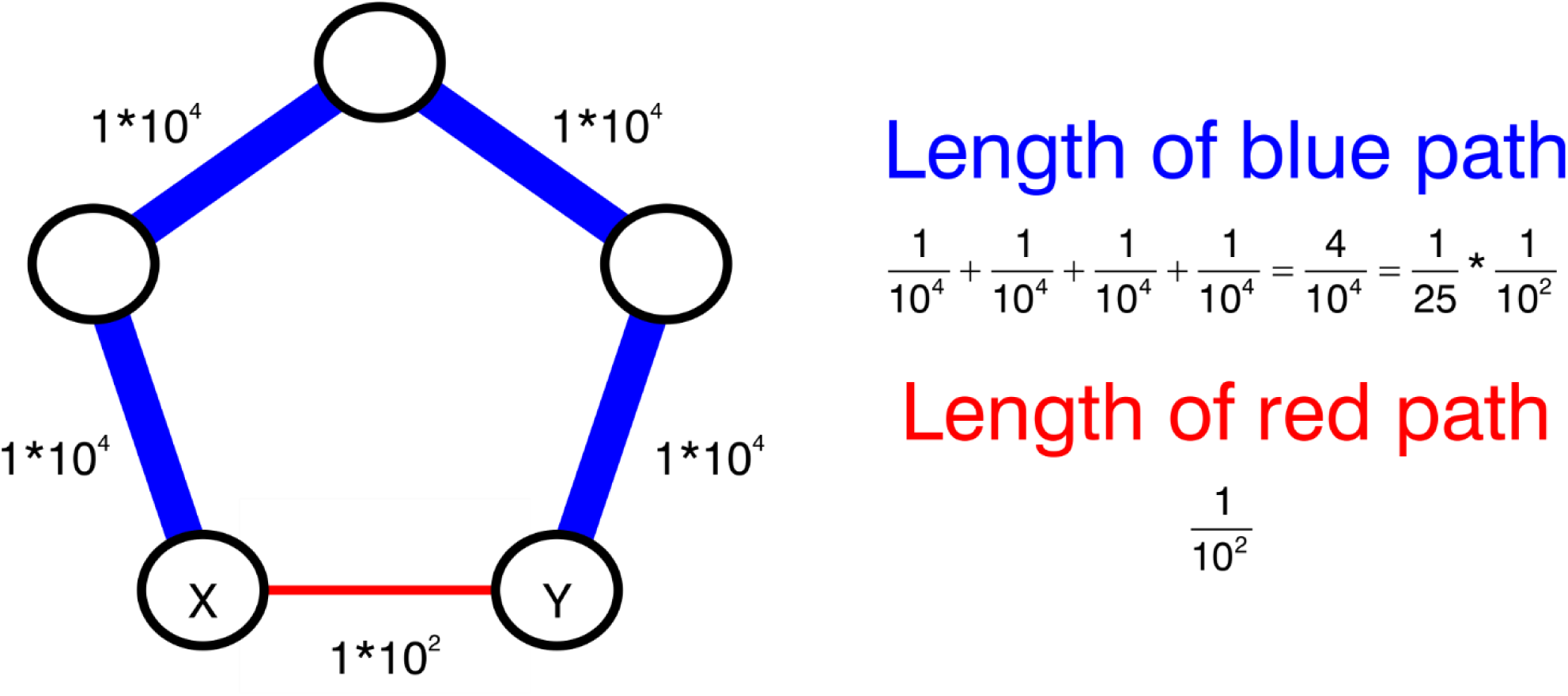
The nature of shortest paths in dense weighted connectomes. Due to the conventional definition of shortest path (the path with the minimum sum of connection ‘lengths’, with length being the reciprocal of connection strength), and the characteristic distribution of dense weighted connectomes (long/heavy tailed), weak connections might be effectively ‘bypassed’ during calculation of metrics that use the shortest path definition. Shown is a toy example illustrating that for traveling between the nodes X and Y, an indirect path made of 4 strong connections (blue) has a lower sum of connection ‘lengths’ than a direct path made of a single weak connection (red). The shortest path between X and Y will be then the blue rather than the red path.

### 4.6. Relationship to findings from studies on weighted structural connectomes

In contrast with our conclusions, the literature includes some claims that weak connections do play an important role when computing graph-theoretical metrics of weighted connectomes. However, the connectomes examined in previous studies differ from our connectomes in at least three important parameters: tracking algorithm, number of streamlines and connectivity density. Drakesmith et al. (2015), for example, evaluated the contribution of weak connections using a connectome generated using deterministic tractography (rather than probabilistic tractography like here), and thus their connectomes were sparse (rather than dense) (for comparison between tractography methods, see Sotiropoulos and Zalesky, 2017). Moreover, the tractogram used for connectome computation was manually edited by removing connections, which made the resulting connectome even sparser. As metrics were shown to be very sensitive to connectivity density in sparse connectomes generated using similar methods (Yan et al., 2018), this is highly likely the reason that the authors found the contribution of even a single weak connection to be statistically significant (given that adding/removing connections obviously affects density). The magnitude of contribution to a metric value by a single weak connection was augmented further due to the relatively small number of streamlines constructed by Drakesmith et al. (resulting in the strongest connection having weight of about 500) compared with the number of streamlines constructed here (strongest connection has weight of ~90,000, see Table 1). Relatively speaking, a connection with strength of 1 in Drakesmith et al.’s study may be equivalent to a connection with strength of about 200 here (in between the 60^th^ and 70^th^ percentile in our typical connection-strength distribution, see Table 1). These methodological differences explain why Drakesmith et al. conclude differently than us that weak connections have a strong adverse effect, and thus must be removed by pruning.

### 4.7. Relationship to findings from studies on functional connectomes

It is often the case that studies on weighted structural connectomes borrow processing methodologies from the literature on functional connectomes. Despite the fact that dense weighted structural connectomes are almost fully connected (and thus resemble weighted functional connectomes which are by definition fully connected), we strongly advocate against it. This is owing to the simple observation that the connection-strength distribution is very different between these two classes of connectomes: long/heavy tail distribution in dense weighted structural connectomes (see Fig. 4) versus approximately Gaussian distribution in weighted functional connectomes (see Fornito et al., 2013). As mentioned above (Section 4.1.1), connection-strength distribution is a major deciding factor in the behaviour of weighted graph-theoretical metrics, and unless explicitly demonstrated, it should not be assumed that connectomes with different connection strength distributions behave the same.

An example of weighted functional connectomes whose connection-strength distribution indeed makes them behave differently than dense weighted connectomes is a class of functional connectomes presented by Ginestet et al. (2011). These functional connectomes are generated by applying a transformation to the original connection-strength distribution, such that the values (i.e. Pearson’s correlation coefficient) in the original range of −1 to 1 are shifted up and rescaled to the range of 0 to 1. This solves the problem of having connections with negative weight, but for typical original distributions (e.g., Fornito et al., 2012), also leads to the weakest connection in the graph being approximately twice as weak as the strongest connection (Ginestet et al., 2011). Ginestet et al. formally showed that, with this type of connection-strength distribution, the shortest path between any two nodes will always be the direct connection between them. No weak connections will be then ‘bypassed’ by stronger ones, and the extreme insensitivity to pruning observed in dense weighted connectomes (see Section 4.5) would not be present.

### 4.8. Limitations

A limitation common to all systematic studies on graph-theoretical metrics is that only a specific set of metrics can be investigated. For our results to be maximally applicable to readers, we focused here on the most popular metrics. We ensured, however, that the metrics chosen are varied enough (e.g., some rely on shortest paths and some do not) to represent a wide range of metric behaviours. It is also of note that we restricted our analysis to the most common definition for each graph-theoretical metric. The only exception is clustering coefficient, where the popular definition proposed by Onnela et al. (2005) was shown to be too strongly dependent on density at high connectivity densities (Yeh et al., 2016). Comparison between different definitions of graph-theoretical metrics is outside the scope of this study.

As with other related studies (Drakesmith et al., 2015; Zalesky et al., 2016), another limitation of our study is that all the graph-theoretical metrics considered are global (i.e. take into account the whole graph and represent its topology using a single number). The investigation of the effect of pruning on metrics that characterize graph topology in a more spatially localized manner (e.g., hubs, rich-club, etc.) warrants future study, as some graph topology traits are known to be poorly captured by the global metrics.

This study is also limited by the fact that tractogram construction and subsequent connectome computation were performed using a single pipeline, with a fixed set of parameters. Our pipeline is considered the state-of-the-art in its class in terms of biological accuracy (Smith et al., 2015a; Yeh et al., 2016), but we cannot rule out that the specific techniques or parameters used had an effect on resulting graph topology, or on its sensitivity to pruning.

## 5. Conclusions

Dense weighted structural connectomes are becoming increasingly common with the use of state-of-the-art fibre tracking methods, but there is no consensus on how to treat the many weak connections that these connectomes include. Our results showed that removal of the weak connections (i.e. pruning) is overall not beneficial; hence, we recommend on the assumption-free no pruning approach. Choosing not to remove weak connections, one avoids an extra analysis step (which can complicate the pipeline) as well as the need to select an arbitrary pruning method and threshold level (or levels). Such arbitrary or heuristic-based choices make studies difficult to compare with each other, and worse, might lead to unstable statistical results at large pruning extents.

## Acknowledgements

We are grateful to the National Health and Medical Research Council (NHMRC) of Australia, the Australian Research Council (ARC), the Victorian Government’s Operational Infrastructure Support Grant, and Melbourne Bioinformatics at the University of Melbourne, grant number UOM0048, for their support. We acknowledge the facilities of the National Imaging Facility, a National Collaborative Research Infrastructure Strategy (NCRIS) capability, at The Florey Institute of Neuroscience and Mental Health. Data were provided in part by the Human Connectome Project, WU-Minn Consortium (Principal Investigators: David Van Essen and Kamil Ugurbil; 1U54MH091657) funded by the 16 NIH Institutes and Centers that support the NIH Blueprint for Neuroscience Research; and by the McDonnell Center for Systems Neuroscience at Washington University. Lastly, we would like to thank Marion Sourty for assistance with HCP data processing, and Donna Parker, Farnoosh Sadeghian, Valerie Yap, Patrick Carney, Magdalena Kowalczyk and Mira Semmelroch for help with subject recruitment and clinical-grade data acquisition.

## Footnotes

1. Given that there is no effective method to date to differentiate between real and spurious connections (Thomas et al., 2014; Maier-Hein et al., 2017), indiscriminate removal of weak connections is used as a common strategy in practice.

2. A preliminary version of this work was presented at the 26th Annual Meeting of the International Society for Magnetic Resonance (Civier et al., 2018).

3. Note that at each target density where the “pruned” and “unpruned” group means cross each other, the *p*-value increases up to 1 and then immediately decreases. Depending on the direction of change that pruning induces on each metric, and on the order between the original means of the “pruned” and “unpruned” groups at each random split, a crossing may occur at least once somewhere along the target density range. Moreover, due to the aforementioned unusual behaviour of the clustering coefficient, for that metric there is often a crossing at 10% density as well.

## References

Andersson, J.L., Skare, S., Ashburner, J., 2003. How to correct susceptibility distortions in spin-echo echo-planar images: application to diffusion tensor imaging. Neuroimage 20, 870–888.

Andersson, J.L., Sotiropoulos, S.N., 2015. Non-parametric representation and prediction of single and multi-shell diffusion-weighted MRI data using Gaussian processes. Neuroimage 122, 166–176.

Andersson, J.L., Sotiropoulos, S.N., 2016. An integrated approach to correction for off-resonance effects and subject movement in diffusion MR imaging. Neuroimage 125, 1063–1078.

Bassett, D.S., Bullmore, E.T., 2016. Small-world brain networks revisited. Neuroscientist 23, 499–516.

Christiaens, D., Reisert, M., Dhollander, T., Sunaert, S., Suetens, P., Maes, F., 2015. Global tractography of multi-shell diffusion-weighted imaging data using a multi-tissue model. Neuroimage 123, 89–101.

Civier, O., Smith, R.E., Yeh, C.-H., Connelly, A., Calamante, F., 2018. Is removal of weak connections necessary for dense weighted structural connectomes?, Joint Annual Meeting ISMRM (International Society for Magnetic Resonance in Medicine) - ESMRMB 2018, Paris, France.

Conti, E., Mitra, J., Calderoni, S., Pannek, K., Shen, K., Pagnozzi, A., Rose, S., Mazzotti, S., Scelfo, D., Tosetti, M., 2017. Network over-connectivity differentiates autism spectrum disorder from other developmental disorders in toddlers: A diffusion MRI study. Human Brain Mapping 38, 2333–2344.

Daducci, A., Canales-Rodríguez, E.J., Descoteaux, M., Garyfallidis, E., Gur, Y., Lin, Y.-C., Mani, M., Merlet, S., Paquette, M., Ramirez-Manzanares, A., 2014. Quantitative comparison of reconstruction methods for intra-voxel fiber recovery from diffusion MRI. IEEE Transactions on Medical Imaging 33, 384–399.

Daducci, A., Gerhard, S., Griffa, A., Lemkaddem, A., Cammoun, L., Gigandet, X., Meuli, R., Hagmann, P., Thiran, J.-P., 2012. The connectome mapper: an open-source processing pipeline to map connectomes with MRI. PloS one 7, e48121.

de Reus, M.A., van den Heuvel, M.P., 2013. Estimating false positives and negatives in brain networks. Neuroimage 70, 402–409.

Desikan, R.S., Segonne, F., Fischl, B., Quinn, B.T., Dickerson, B.C., Blacker, D., Buckner, R.L., Dale, A.M., Maguire, R.P., Hyman, B.T., Albert, M.S., Killiany, R.J., 2006. An automated labeling system for subdividing the human cerebral cortex on MRI scans into gyral based regions of interest. Neuroimage 31, 968–980.

Drakesmith, M., Caeyenberghs, K., Dutt, A., Lewis, G., David, A.S., Jones, D.K., 2015. Overcoming the effects of false positives and threshold bias in graph theoretical analyses of neuroimaging data. Neuroimage 118, 313–333.

Ercsey-Ravasz, M., Markov, N.T., Lamy, C., Van Essen, D.C., Knoblauch, K., Toroczkai, Z., Kennedy, H., 2013. A predictive network model of cerebral cortical connectivity based on a distance rule. Neuron 80, 184–197.

Feinberg, D.A., Moeller, S., Smith, S.M., Auerbach, E., Ramanna, S., Glasser, M.F., Miller, K.L., Ugurbil, K., Yacoub, E., 2010. Multiplexed echo planar imaging for sub-second whole brain FMRI and fast diffusion imaging. PloS one 5, e15710.

Fornito, A., Zalesky, A., Breakspear, M., 2013. Graph analysis of the human connectome: promise, progress, and pitfalls. Neuroimage 80, 426–444.

Fornito, A., Zalesky, A., Pantelis, C., Bullmore, E.T., 2012. Schizophrenia, neuroimaging and connectomics. Neuroimage 62, 2296–2314.

Ginestet, C.E., Nichols, T.E., Bullmore, E.T., Simmons, A., 2011. Brain network analysis: separating cost from topology using cost-integration. PloS one 6, e21570.

Girard, G., Daducci, A., Petit, L., Thiran, J.P., Whittingstall, K., Deriche, R., Wassermann, D., Descoteaux, M., 2017. Ax Tract: Toward microstructure informed tractography. Human Brain Mapping 38, 5485–5500.

Gorgolewski, K., Burns, C.D., Madison, C., Clark, D., Halchenko, Y.O., Waskom, M.L., Ghosh, S.S., 2011. Nipype: a flexible, lightweight and extensible neuroimaging data processing framework in python. Frontiers in neuroinformatics 5, 13.

Hagmann, P., Cammoun, L., Gigandet, X., Meuli, R., Honey, C.J., Wedeen, V.J., Sporns, O., 2008. Mapping the structural core of human cerebral cortex. PLoS biology 6, e159.

Hagmann, P., Kurant, M., Gigandet, X., Thiran, P., Wedeen, V.J., Meuli, R., Thiran, J.-P., 2007. Mapping human whole-brain structural networks with diffusion MRI. PloS one 2, e597.

Iturria-Medina, Y., Sotero, R.C., Canales-Rodríguez, E.J., Alemán-Gómez, Y., Melie-García, L., 2008. Studying the human brain anatomical network via diffusion-weighted MRI and graph theory. Neuroimage 40, 1064–1076.

Jeurissen, B., Tournier, J.-D., Dhollander, T., Connelly, A., Sijbers, J., 2014. Multi-tissue constrained spherical deconvolution for improved analysis of multi-shell diffusion MRI data. Neuroimage 103, 411–426.

Jones, D.K., Knosche, T.R., Turner, R., 2013. White matter integrity, fiber count, and other fallacies: The do’s and don’ts of diffusion MRI. Neuroimage 73, 239–254.

Kamagata, K., Zalesky, A., Hatano, T., Di Biase, M.A., El Samad, O., Saiki, S., Shimoji, K., Kumamaru, K.K., Kamiya, K., Hori, M., 2018. Connectome analysis with diffusion MRI in idiopathic Parkinson’s disease: Evaluation using multi-shell, multi-tissue, constrained spherical deconvolution. NeuroImage: Clinical 17, 518–529.

Kellner, E., Dhital, B., Kiselev, V.G., Reisert, M., 2016. Gibbs-ringing artifact removal based on local subvoxel-shifts. Magnetic Resonance in Medicine 76, 1574–1581.

Lemkaddem, A., Skiöldebrand, D., Dal Palú, A., Thiran, J.-P., Daducci, A., 2014. Global tractography with embedded anatomical priors for quantitative connectivity analysis. Frontiers in neurology 5, 232.

Markov, N., Misery, P., Falchier, A., Lamy, C., Vezoli, J., Quilodran, R., Gariel, M., Giroud, P., Ercsey-Ravasz, M., Pilaz, L., 2011. Weight consistency specifies regularities of macaque cortical networks. Cerebral Cortex 21, 1254–1272.

Moeller, S., Yacoub, E., Olman, C.A., Auerbach, E., Strupp, J., Harel, N., Uğurbil, K., 2010. Multiband multislice GE-EPI at 7 tesla, with 16-fold acceleration using partial parallel imaging with application to high spatial and temporal whole-brain fMRI. Magnetic Resonance in Medicine 63, 1144–1153.

Mugler III, J.P., Brookeman, J.R., 1990. Three-dimensional magnetization-prepared rapid gradient-echo imaging (3D MP RAGE). Magnetic Resonance in Medicine 15, 152–157.

Oh, S.W., Harris, J.A., Ng, L., Winslow, B., Cain, N., Mihalas, S., Wang, Q., Lau, C., Kuan, L., Henry, A.M., 2014. A mesoscale connectome of the mouse brain. Nature 508, 207.

Onnela, J.-P., Saramäki, J., Kertész, J., Kaski, K., 2005. Intensity and coherence of motifs in weighted complex networks. Physical Review E 71, 065103.

Oxtoby, N.P., Garbarino, S., Firth, N.C., Warren, J.D., Schott, J.M., Alexander, D.C., Initiative, A.s.D.N., 2017. Data-driven sequence of changes to anatomical brain connectivity in sporadic Alzheimer’s disease. Frontiers in neurology 8, 580.

Patenaude, B., Smith, S.M., Kennedy, D.N., Jenkinson, M., 2011. A Bayesian model of shape and appearance for subcortical brain segmentation. Neuroimage 56, 907–922.

Perry, A., Wen, W., Lord, A., Thalamuthu, A., Roberts, G., Mitchell, P.B., Sachdev, P.S., Breakspear, M., 2015. The organisation of the elderly connectome. Neuroimage 114, 414–426.

Pestilli, F., Yeatman, J.D., Rokem, A., Kay, K.N., Wandell, B.A., 2014. Evaluation and statistical inference for human connectomes. Nature methods 11, 1058.

Reese, T.G., Heid, O., Weisskoff, R., Wedeen, V., 2003. Reduction of eddy-current-induced distortion in diffusion MRI using a twice-refocused spin echo. Magnetic Resonance in Medicine 49, 177–182.

Reisert, M., Mader, I., Anastasopoulos, C., Weigel, M., Schnell, S., Kiselev, V., 2011. Global fiber reconstruction becomes practical. Neuroimage 54, 955–962.

Rubinov, M., Sporns, O., 2010. Complex network measures of brain connectivity: uses and interpretations. Neuroimage 52, 1059–1069.

Rubinov, M., Sporns, O., 2011. Weight-conserving characterization of complex functional brain networks. Neuroimage 56, 2068–2079.

Setsompop, K., Gagoski, B.A., Polimeni, J.R., Witzel, T., Wedeen, V.J., Wald, L.L., 2012. Blipped-controlled aliasing in parallel imaging for simultaneous multislice echo planar imaging with reduced g-factor penalty. Magnetic Resonance in Medicine 67, 1210–1224.

Sherbondy, A.J., Rowe, M.C., Alexander, D.C., 2010. MicroTrack: an algorithm for concurrent projectome and microstructure estimation. International Conference on Medical Image Computing and Computer-Assisted Intervention. Springer, pp. 183–190.

Smith, R.E., Tournier, J.-D., Calamante, F., Connelly, A., 2012. Anatomically-constrained tractography: improved diffusion MRI streamlines tractography through effective use of anatomical information. Neuroimage 62, 1924–1938.

Smith, R.E., Tournier, J.-D., Calamante, F., Connelly, A., 2013. SIFT: spherical-deconvolution informed filtering of tractograms. Neuroimage 67, 298–312.

Smith, R.E., Tournier, J.-D., Calamante, F., Connelly, A., 2015a. The effects of SIFT on the reproducibility and biological accuracy of the structural connectome. Neuroimage 104, 253–265.

Smith, R.E., Tournier, J.-D., Calamante, F., Connelly, A., 2015b. SIFT2: Enabling dense quantitative assessment of brain white matter connectivity using streamlines tractography. Neuroimage 119, 338–351.

Smith, S.M., Jenkinson, M., Woolrich, M.W., Beckmann, C.F., Behrens, T.E., Johansen-Berg, H., Bannister, P.R., De Luca, M., Drobnjak, I., Flitney, D.E., 2004. Advances in functional and structural MR image analysis and implementation as FSL. Neuroimage 23, S208–S219.

Sotiropoulos, S., Moeller, S., Jbabdi, S., Xu, J., Andersson, J., Auerbach, E., Yacoub, E., Feinberg, D., Setsompop, K., Wald, L., 2013. Effects of image reconstruction on fiber orientation mapping from multichannel diffusion MRI: reducing the noise floor using SENSE. Magnetic Resonance in Medicine 70, 1682–1689.

Sotiropoulos, S.N., Zalesky, A., 2017. Building connectomes using diffusion MRI: why, how and but. NMR in Biomedicine. https://doi.org/10.1002/nbm.3752

Sporns, O., Tononi, G., Kotter, R., 2005. The human connectome: A structural description of the human brain. PLoS Comput Biol 1, e42.

Tournier, J.-D., Calamante, F., Connelly, A., 2007. Robust determination of the fibre orientation distribution in diffusion MRI: non-negativity constrained super-resolved spherical deconvolution. Neuroimage 35, 1459–1472.

Tournier, J.-D., Calamante, F., Gadian, D.G., Connelly, A., 2004. Direct estimation of the fiber orientation density function from diffusion-weighted MRI data using spherical deconvolution. Neuroimage 23, 1176–1185.

Tournier, J.D., Calamante, F., Connelly, A., 2010. Improved probabilistic streamlines tractography by 2^nd^ order integration over fibre orientation distributions. Proceedings of the international society for magnetic resonance in medicine, p. 1670.

Tournier, J.D., Mori, S., Leemans, A., 2011. Diffusion tensor imaging and beyond. Magnetic Resonance in Medicine 65, 1532–1556.

Tustison, N.J., Avants, B.B., Cook, P.A., Zheng, Y., Egan, A., Yushkevich, P.A., Gee, J.C., 2010. N4ITK: improved N3 bias correction. IEEE Transactions on Medical Imaging 29, 1310–1320.

Van Essen, D.C., Smith, S.M., Barch, D.M., Behrens, T.E., Yacoub, E., Ugurbil, K., Consortium, W.- M.H., 2013. The WU-Minn human connectome project: an overview. Neuroimage 80, 62–79.

van Wijk, B.C., Stam, C.J., Daffertshofer, A., 2010. Comparing brain networks of different size and connectivity density using graph theory. PLoS One 5, e13701.

Vasa, F., Seidlitz, J., Romero-Garcia, R., Whitaker, K.J., Rosenthal, G., Vertes, P.E., Shinn, M., Alexander-Bloch, A., Fonagy, P., Dolan, R.J., Jones, P.B., Goodyer, I.M., consortium, N., Sporns, O., Bullmore, E.T., 2018. Adolescent tuning of association cortex in human structural brain networks. Cerebral Cortex 28, 281–294.

Veraart, J., Fieremans, E., Novikov, D.S., 2016a. Diffusion MRI noise mapping using random matrix theory. Magnetic Resonance in Medicine 76, 1582–1593.

Veraart, J., Novikov, D.S., Christiaens, D., Ades-Aron, B., Sijbers, J., Fieremans, E., 2016b. Denoising of diffusion MRI using random matrix theory. Neuroimage 142, 394–406.

Xu, J., Moeller, S., Strupp, J., Auerbach, E., Chen, L., Feinberg, D., Ugurbil, K., Yacoub, E., 2012. Highly accelerated whole brain imaging using aligned-blipped-controlled-aliasing multiband EPI. Proceedings of the 20^th^ Annual Meeting of ISMRM.

Yan, X., Jeub, L.G., Flammini, A., Radicchi, F., Fortunato, S., 2018. Weight thresholding on complex networks. Physical Review E. arXiv preprint arXiv:1806.07479.

Yeh, C.H., Smith, R.E., Liang, X., Calamante, F., Connelly, A., 2016. Correction for diffusion MRI fibre tracking biases: The consequences for structural connectomic metrics. Neuroimage 142, 150–162.

Ypma, R.J., Bullmore, E.T., 2016. Statistical analysis of tract-tracing experiments demonstrates a dense, complex cortical network in the mouse. PLoS computational biology 12, e1005104.

Zalesky, A., Fornito, A., Cocchi, L., Gollo, L.L., van den Heuvel, M.P., Breakspear, M., 2016. Connectome sensitivity or specificity: which is more important? Neuroimage 142, 407–420.

Zhang, B., Horvath, S., 2005. A general framework for weighted gene co-expression network analysis. Statistical applications in genetics and molecular biology 4.

