## Supplementary Figures for "Is removal of weak connections necessary for graph-theoretical analysis of dense weighted structural connectomes?"

### Supplementary material S1

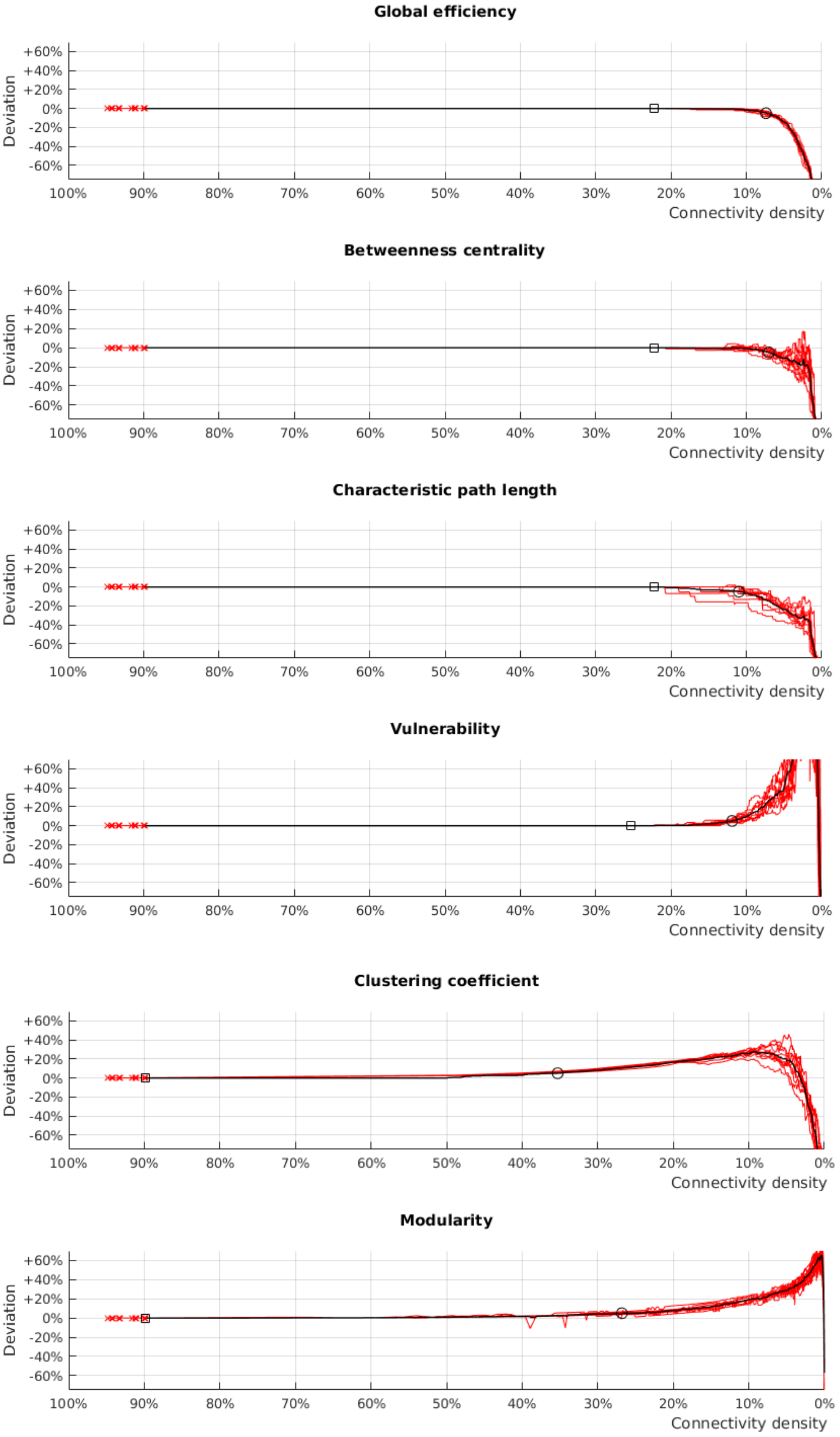


**Fig. S1. The effect of pruning on graph-theoretical metrics (weighted versions) of dense weighted structural connectomes computed from clinical scanner-grade data.** Shown are pruning-induced changes for 10 individuals, as well as changes in their group mean. Conventions are as in Fig. 1.

### Supplementary material S2

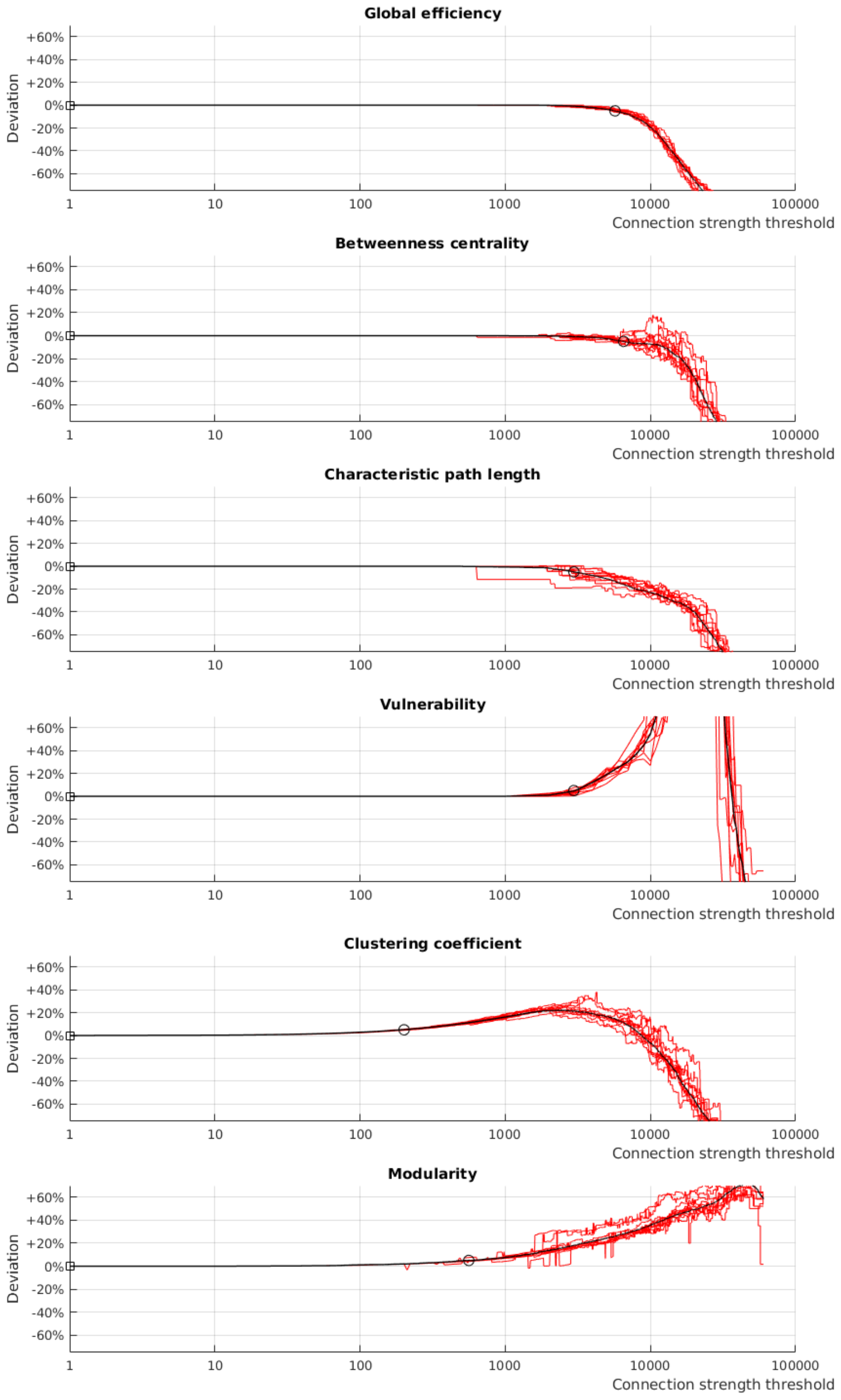


**Fig. S2. The effect of pruning by connection strength threshold on graph-theoretical metrics (weighted versions) of dense weighted structural connectomes.** Shown are pruning-induced changes for 10 subjects from the HCP cohort, as well as changes in their group mean. Conventions are as in Fig. 1, other than x-axis which here specifies threshold values for connection strength (logarithmic scale). For comparison with the x-axis of Fig. 1, please refer to Table 1. For example, pruning to a threshold value of 3009 ($90^{th}$ percentile of the connection strength distribution) is roughly equivalent to target density of 100 – 90 = 10%.

### Supplementary material S3

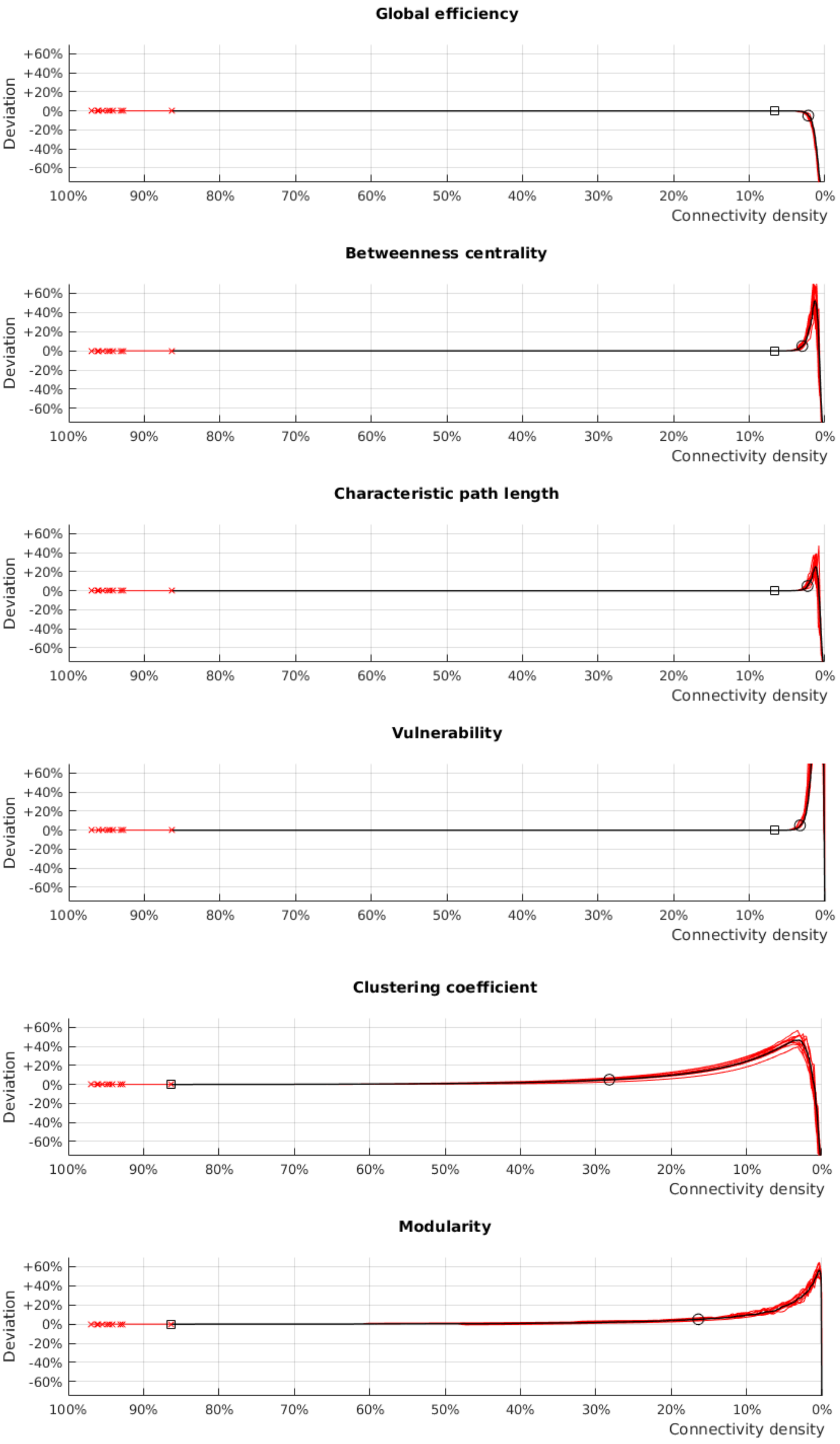


**Fig. S3. The effect of pruning on graph-theoretical metrics (weighted versions) of dense weighted structural connectomes computed using the Laussanne2008 parcellation (234 regions version).** Shown are pruning-induced changes for 10 subjects from the HCP cohort, as well as changes in their group mean. Conventions are as in Fig. 1.
